# Competitive learning suggests circulating miRNA profiles for cancers decades prior to diagnosis

**DOI:** 10.1101/2020.03.26.009597

**Authors:** Andreas Keller, Tobias Fehlmann, Christina Backes, Fabian Kern, Randi Gislefoss, Hilde Langseth, Trine B. Rounge, Nicole Ludwig, Eckart Meese

## Abstract

Small non-coding RNAs such as microRNAs are master regulators of gene expression. One of the most promising applications of miRNAs is the use as liquid biopsy. Especially early diagnosis is an effective means to increase patients’ overall survival. E.g. in oncology a tumor is detected at best prior to its clinical manifestation. We generated genome-wide miRNA profiles from serum of patients and controls from the population-based Janus Serum Bank (JSB) and analyzed them by bioinformatics and artificial intelligence approaches. JSB contains sera from 318,628 originally healthy persons, more than 96,000 of whom later developed cancer. We selected 210 serum samples of patients with lung, colon or breast cancer at three time points prior to diagnosis, after cancer diagnosis and controls. The controls were matched with regard to age of the blood donor and to the time points of blood drawing, which were 27, 32, or 38 years prior to diagnosis. Using ANOVA we report 70 significantly deregulated markers (adjusted p-value<0.05). The driver for the significance was the diagnostic time point (miR-575, miR-6821-5p, miR-630 had adjusted p-values<10^−10^). Further, 91miRNAs were differently expressed in pre-diagnostic samples as compared to controls (nominal p<0.05). Unsupervised competitive learning by self-organized maps indicated larges effects in lung cancer samples while breast cancer samples showed the least pronounced changes. Self-organized maps also highlighted cancer and time point specific miRNA dys-regulation. Intriguingly, a detailed breakdown of the results highlighted that 51% of all miRNAs were highly specific, either for a time-point or a cancer entity. Our results indicate that tumors may be indicated by serum miRNAs decades prior the clinical manifestation.

## Introduction

MicroRNAs – often abbreviated as miRs or miRNAs – are short non-coding RNAs. In basically all organisms, miRNAs regulate the gene expression on posttranscriptional level by binding to the 3’UTR of target genes^1, 2^. MiRNAs control various cellular processes by targeting a broad number of different target genes and target pathways. Even non-canonical binding sites as short as 5-mers can have a deterministic influence on the targeting process^3^. Many diseases including various cancer types as well as neurodegenerative diseases are associated with aberrant miRNA expression in affected tissues and body fluids^4–7^. One of the most promising applications of miRNAs is to facilitate early disease detection as liquid biopsies, especially in cancer.

Cancer is the second-most common cause of death worldwide and the most common cause of death in men and women under the age of 70 and has become a large public health problem. The challenges in cancer management are to succeed in early detection, to improve diagnostic precision, to offer an appropriate therapy and follow-up, all aimed at reducing suffering and prolonging survival. Early detection is especially important regarding reduced mortality as therapeutic intervention of advanced cancers often has low effect on survival. Early cancer detection is particularly promising if the tumor detection occurs prior to the clinical manifestation. There is, however, a paucity of studies that pursue the latter idea due to the lack of prospectively collected biospecimens. We used samples of the Janus Serum Bank, a unique population-based biobank, which collected sera form 318,628 originally healthy persons, over 96,000 of whom developed cancer after the first samples were taken ^8–11^. One advantage of the Janus Serum bank is the multiple and regular sampling over time allowing to follow up pre-diagnostic changes of a biomarker.

In addition to miRNAs, the stored serum contains other molecules which can be tested for their predictive potential to indicate tumor development at very early stages^12^. Examples in sum include proteins, DNA, metabolites, small non-coding RNAs and epigenetic changes^13–15^. Especially, their high degree of stability in blood has driven the search for miRNAs biomarkers. As for any biomarker the identification of confounding factors is essential to estimate the diagnostic value of blood-borne miRNAs^6, 16^. Previously, we comprehensively evaluated the influence of storage time on the totality of blood-borne miRNAs by analyzing consecutive samples of healthy individuals that have been stored in the Janus Serum Bank between 23 and 40 years at −25 °Celsius. We found that a substantial proportion of the miRNome was affected by the age of the blood donor but only few miRNAs showed variations in their abundance depending on their storage time (measurement)^17^. Also other factors such as smoking influenced the miRNA expression significantly^18^. Furthermore, we previously also provided first and preliminary indications for pre-diagnostic miRNA profiles^19^ in serum of individuals, who were later diagnosed with lung cancer^20^.

Because of the urgent clinical needs for screening markers, much research is dedicated to discover molecular pre-diagnostic cancer signatures. For example, DNA methylation changes measured in prediagnostic peripheral blood samples were found to be associated with smoking and lung cancer risk^21^. In a similar direction, hypomethylation of smoking-related genes was observed to be correlated to future lung cancer in four prospective cohorts^22^. Further, pre-diagnostic leukocyte mitochondrial DNA copy number has been discussed in the context of lung cancer risk^23^. Also autoantibodies against tumor-associated antigens seem to have a potential for positive results in pre-diagnostic samples^24^. For colon cancer a meta-analysis investigated pre-diagnostic protein levels. The results of the study suggest an association of pre-diagnostic circulating CRP levels with an increased risk of colorectal cancer ^25^. Similarly, pre-diagnostic levels of adiponectin and soluble vascular cell adhesion molecule-1 (VCAM1) seem to be associated with colorectal cancer risk^26^. For breast cancer, biomarker candidates have been likewise identified using serum protein profiling of pre-diagnostic serum^27^. But also in other cancer types, such as ovarian cancer, pre-diagnostic signatures were identified^28^. The list of other studies on pre-diagnostic biomarkers is far from being complete, however, it show the potential of respective tests and the high research interest in pre-diagnostic cancer markers.

These previous studies often rely on one or few markers. There is however a clear trend towards more complex biomarker sets. Further, biomarker studies now often consider more than only one disease at a time^14^. To analyze respective complex studies including multiple time points and multiple cancer types different bioinformatics, biostatistics, machine learning or artificial intelligence approaches can be applied. Given the nature of the study we avoided to use supervised learning but tested unsupervised competitive learning approaches. Respective approaches such as Self Organized Maps (SOMs), originally introduced in 1982 by Kohonen^29^ support the discovery of structures in the data. SOMs typically generate twodimensional and well interpretable discretized representations of a high dimensional input space. Using SOMs and classical biostatistics methods we set to extend the knowledge on pre-diagnostic miRNA biomarkers in serum by including strictly matched controls and by analyzing samples from patients with carcinoma of lung, colon and breast. We address the questions if and how long prior to the diagnosis characteristic blood-borne miRNA changes can be observed in these cases and whether specific pre-diagnostic miRNA signatures can be found for these cancer types. We start our consideration with global aspects, i.e. we try to identify overall pre-diagnostic cancer markers before we address the topic of discovering prediagnostic markers that are specific for one cancer entity.

## Results

### Study Set up and miRNA profiling

We selected serum samples of individuals that later developed cancer out of 318,628 stored samples of the Janus Serum Bank. The individuals had samples at three time points, with five-six year intervals, prior to cancer diagnosis and one time point after diagnosis for each patient. Stringent selected cancer-free controls with the following criteria: i) the matched controls stemmed from donors of the same sex as the cases, ii) the age difference between cases and matched controls was not more than two years, and iii) the difference between the blood collection time point of cases and controls was not more than two months ensured control for confounding factors. The cases had to develop either lung, breast or colon cancer but were not diagnosed with any other cancer type prior to diagnosis. The included matched controls were not diagnosed with cancer at any time (Fig. 1a/b). Based on these criteria we were able to identify 90 case-control paired samples. The samples were stored for up to 40 years with a median storage time of 33 years and a median age of blood donors of 41 years at enrolment (Fig. 1c). Most samples and cases were collected at three time points i.e. at 27, 32, or 38 years prior to diagnosis (Fig. 1d). Inherent to the longitudinal character of the study design is a strong negative correlation of the age of donors and the storage length, i.e. the earliest samples stem necessarily from the youngest cases and controls (Fig. 1d). In total, we analyzed the miRNomes of 210 samples including 120 samples from cases and 90 from controls. Six samples (2.9%) yielded miRNomes of low quality and were excluded from further analysis. Of 2,549 profiled miRNAs, 435 were expressed above the background in the serum samples. Principle component analysis highlighted the collection time point as major contributing factor to the variance in the miRNA data emphasizing the importance of the stringent study design with closely matched controls (Fig. 1e).

**Figure 1:**
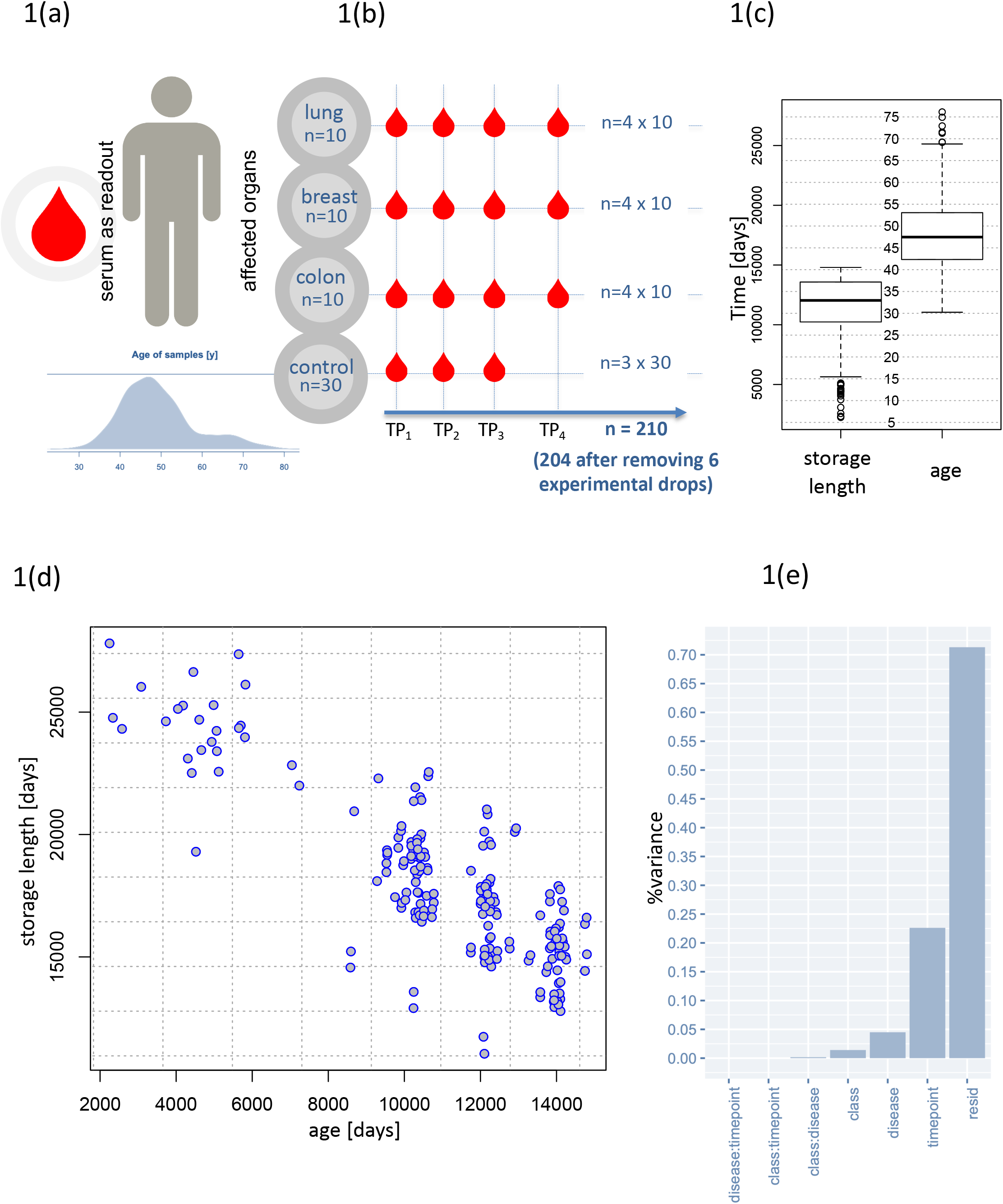
Study set-up and characteristics of the participants. (a) Age distribution and read out. (b) Sampling by cancer type and time point (TP). TP1, 2 and 3 refer to prediagnostic sampling time points, TP4 is a sampling time-point after cancer diagnosis. (c) Distribution of storage length and age of blood donors (in days and years) for all samples. (d) Correlation between storage length (in days) and time-points of blood collection (in days). The grouping of the three vertical clusters reflects the fact that most collection time points were 27 years, 32 years and 38 years prior to diagnosis. (e) Percent of data variance for the disease (class; cancer vs controls) the affected organ (lung, colon, breast) and the time point (TP1, TP2, TP3, TP4).

### General patterns between controls, cancer patients prior to and post diagnosis

First, we assessed differences between the control samples, the pre-diagnostic and the postdiagnostic samples without considering neither the cancer type nor the longitudinal aspect, i.e. the three collection time points. An analysis of variance (ANOVA) for all cancer types together and with the three groups (controls, pre-diagnostic and post-diagnostic) identified 134 significantly deregulated miRNAs (unadjusted p < 0.05). Following adjustment using the Benjaimin-Hochberg approach still 70 markers remained significant at an alpha level of 0.05. The three most significant miRNAs (miR-575, miR-6821-5p, and miR-630) showed adjusted p-values of below 10^-10^. For each miRNA, raw and adjusted p-value are detailed in Supplemental Table 1. For the majority of miRNAs, the controls largely matched to the samples collected prior to diagnosis. In contrast, the post-diagnostic cancer showed significantly different miRNA expression as compared to the controls. Examples are provided in Fig. 2a/b showing miRNAs with either reduced or elevated expression in the postdiagnostic samples as compared to similar expression between controls and pre-diagnostic samples. This calls for a more specific analysis in order to discover miRNAs that are differently expressed between controls and pre-diagnostic samples. As shown by the density distribution of the p-values, there are highly significant miRNAs found for the comparison of pre-diagnostic samples and controls (Fig. 2c). Wilcoxon Mann-Whitney (WMW) tests for these two groups identified 91 significant differently expressed miRNAs (Supplemental Table 2; Fig. 2c). We next computed the AUC for each of these miRNAs and identified a bi-variate distribution with one peak representing miRNA with lower expression in the pre-diagnostic cancer samples and the other peak representing miRNAs with higher expression in the prediagnostic samples, each as compared to the controls. The AUC analysis shows that the number of the overexpressed miRNAs is lower (40 miRNAs) than the number of the miRNAs with lower expression (51 miRNAs) in the pre-diagnostic samples as compared to the controls (Fig. 2d). Fig. 2e shows miR-149-3p as an example of a miRNA with higher expression in the pre-diagnostic cases than in the controls.

**Figure 2:**
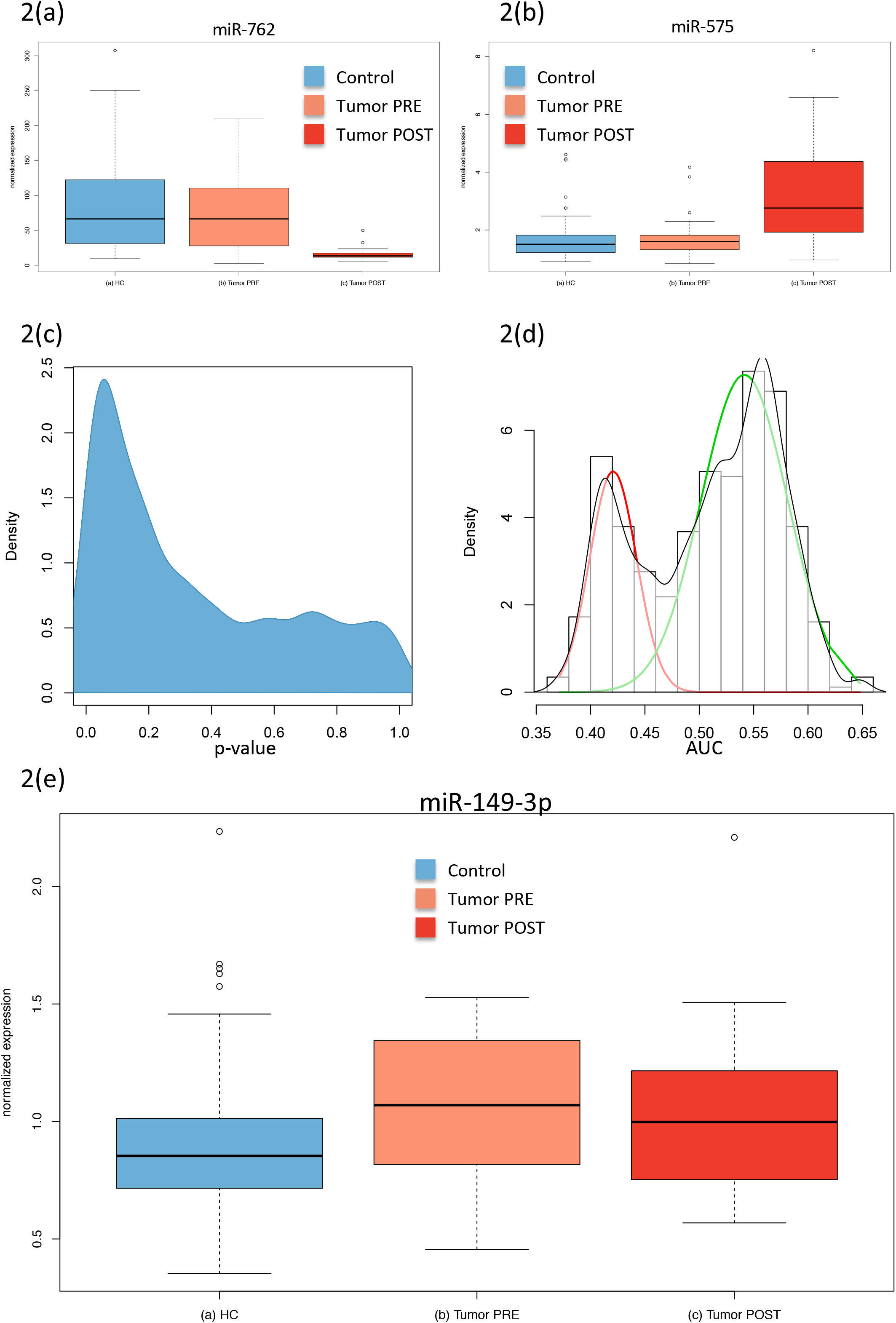
Comparison between cancer and control samples. (a) Box-plot of the normalized expression values of miR-762 in all control samples (free from cancer), all samples collected prior to cancer diagnosis (Cancer PREsampling) and all samples collected after cancer diagnosis (Cancer POSTsampling). (b) Box-plot of the normalized expression values of miR-575 according to Figure 2a. (c) Density distribution of unadjusted p-values for the comparison between all pre sampling time points and the matched controls, showing an enrichment of low (significant) p-values. (d) AUC distributions for the comparison between all cancer pre sampling time points and the matched controls. An AUC close to 1 indicates higher expression in cancer samples and an AUC close to 0 indicate higher expression in the control samples. The red curve corresponds to miRNAs with higher expression in the control samples and the green curve to miRNAs with higher expression in pre-diagnostic cancer samples. (e) Box-plot of the normalized expression values for miR-149-3p showing higher expression in pre-diagnostic cancer samples compared to the matched controls. The labeling is as in Fig. 2a.

### Artificial intelligence to learn temporal and disease specific patterns

We next included in our analysis both the cancer type and the temporal aspect, i.e. the three collection time points. To this end, we applied the self-organizing map (SOM) as a competitive learning based artificial neural network. SOM is an unsupervised approach, but we used the AUC (and with this also the cancer / control information) as input. The SOM facilitated dimension reduction, bringing the 435-dimensional miRNA space to a 2-dimensional representation. First, the SOM was trained with data from all cancer cases combining pre- and postdiagnostic samples as compared to the control samples. In general, the SOM identified three major groups of deregulated miRNAs in this comparison including miRNAs with lower expression in the cancer samples, unaffected miRNAs and miRNAs with higher expression in the cancer cases (Fig. 3a). Second, SOM was trained with data from the three pre-diagnostic time points each compared to the control samples. Throughout all three time points the SOMs reveal a group of miRNAs with lower expression in pre-diagnostic cancer samples as compared to the controls (Fig. 3b, c, d). Higher expressed miRNAs in pre-diagnostic cancer samples were only identified for the time points closest to diagnosis (Fig. 3d). The comparison between the post-diagnostic samples and the controls highlights distinct patterns of both lower and higher expressed miRNAs in the cancer samples (Fig. 3e). In summary, SOM showed that the general pattern trained with data from all cancer cases combining and all control samples (Fig. 3a) is a composition of the disjointed sub-patterns of the prediagnostic and of the post-diagnostic miRNAs.

**Figure 3:**
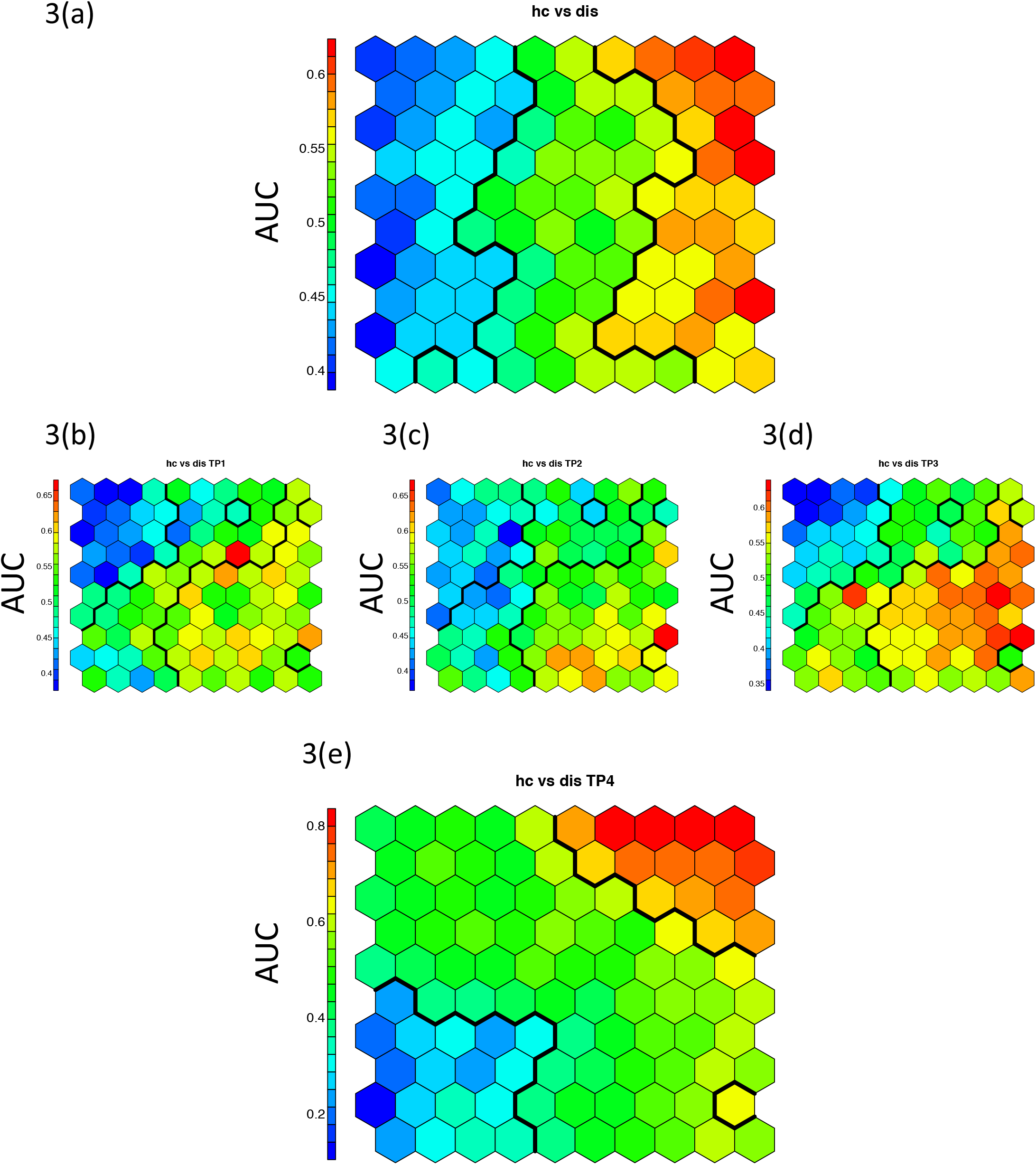
Self-organized maps (SOMs) independent of cancer type. The SOMs were trained with data from all cancer cases and all control samples. Each hexagon represents at least one but usually a set of miRNAs. The color of a hexagon represents the AUC values of the miRNAs in that hexagon with the color code indicated to the left of each subfigure. (a) SOM for the comparison between all samples of the cancer patients and all control samples. Hierarchical clustering identified vertical sections indicating three clusters of differentially expressed miRNAs. (b) SOM for the comparison between of all samples collected at the first time point prior to cancer diagnosis (TimePoint1) as compared to matched controls. There is a cluster of miRNAs with lower expression in the pre-diagnostic cancer samples (indicated in blue in the upper left corner). (c) SOM for the comparison between of all samples collected at the second time point prior to cancer diagnosis (TimePoint2) as compared to matched controls. The cluster of miRNAs with lower expression in pre-diagnostic cancer samples is less evident than for the first time point. (d) SOM for the comparison between of all samples collected at the third time point prior to cancer diagnosis (TimePoint3) as compared to matched controls. There is again a cluster of miRNAs with lower expression but also a cluster with higher expression in pre-diagnostic cancer samples. (e) SOM for the comparison between of all samples collected after cancer diagnosis (Time Point 4) as compared to combined controls (**h**ealthy **c**ontrols). There is again a cluster of miRNAs with lower expression (indicated in blue in the lower left corner) and a cluster with higher expression (indicated in red/orange in the upper left corner) in post-diagnostic cancer samples. Notable, the overall SOM in Fig. 3a largely comprises both, the blue and red clusters from the comparisons shown in Fig. 3b-e.

We next investigated separately the patterns for the three cancer types lung, colon and breast. Specifically, the SOMs were analyzed for each cancer type according to the following five scenarios: All cancer samples versus matched controls, the pre-cancer samples of the first, second, and third time point each compared to matched controls, and the post-diagnostic samples compared to the combined controls. The resulting 3 x 5 patterns are shown in Fig. 4. The analysis of the samples combined for each cancer type identified three groups of expressed miRNAs including lower expressed miRNAs, unaffected miRNAs and higher expressed miRNAs for all three cancer types.

**Figure 4.**
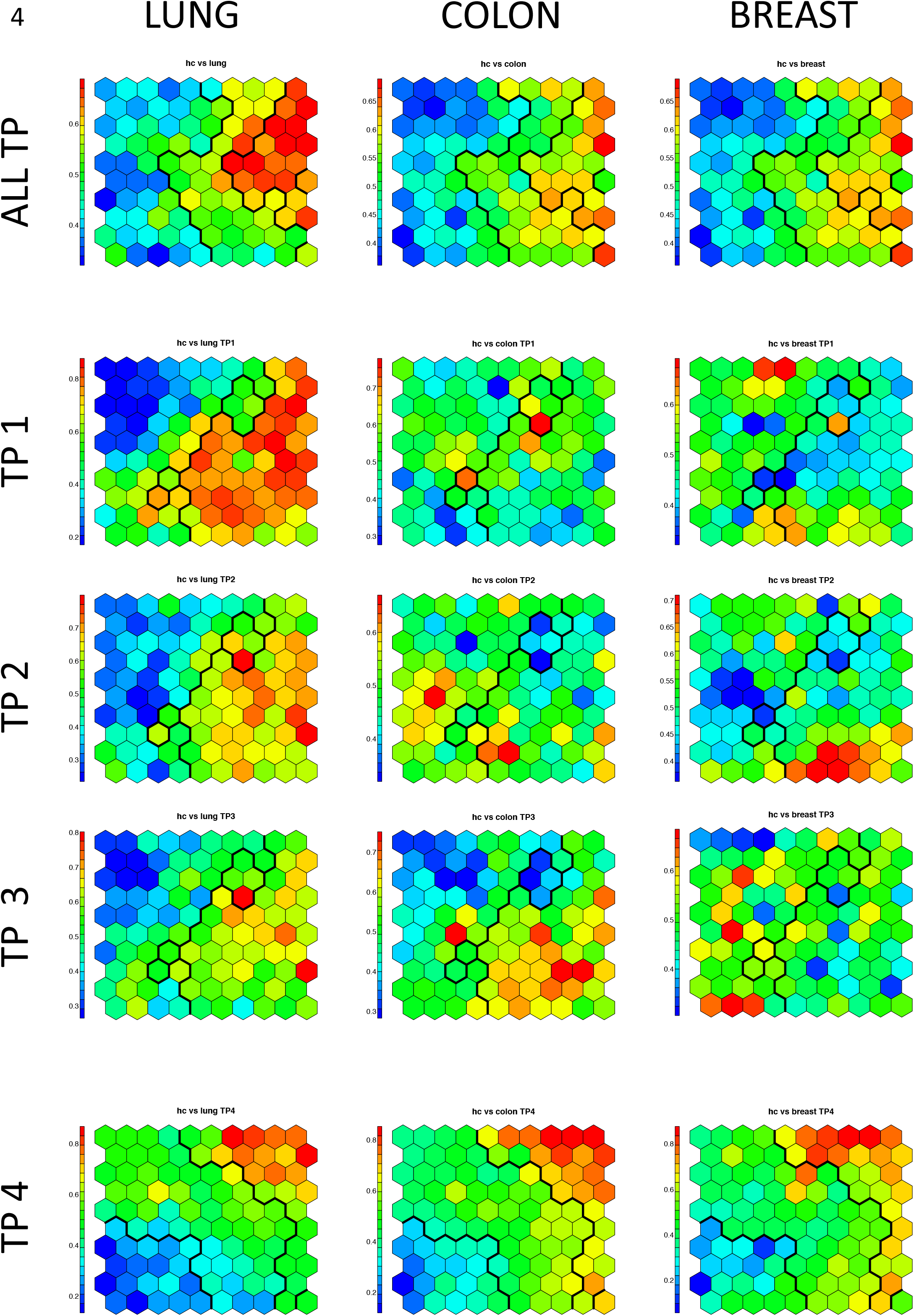
Self-organized maps (SOMS) considering the tumor type. The experimental set-up and the Figure lay-out were as in Fig. 3. The analysis was done for each tumor type separately. The tumor types are indicated on top and the comparisons to the left of the figure (**All T**ime**P**oints; **T**ime**P**oint 1, 2, 3, 4). The most prominent differences between the three tumor types are found for time points 1, 2, and 3. There is a cluster of miRNAs with lower expression in samples collected prior to the diagnosis of lung cancer for all three time points. A cluster of miRNAs with lower expression in samples collected prior to the diagnosis of colon cancer occurs only at time point 3, i.e. closest to diagnosis. There is no comparable clustering for breast cancer patients.

The comparison between the three pre-diagnostic time points and the matched controls revealed specific patterns for each of the three cancer types. The SOM revealed a group of up-regulated miRNAs for the first pre-diagnostic time point in lung cancer samples. This group of up-regulated miRNAs was not found at the second or third time points for lung cancer. In contrast, the SOM did not describe clear signatures for colon and breast cancer at the first and second time points. For colon cancer, the SOM identified a group of down-regulated miRNAs at time point three similar to the pattern observed for lung cancer at time point three. The SOM did not reveal clear patterns of up- or down-regulated miRNAs for breast cancer.

The SOM revealed distinct patterns for each cancer type for the comparison between postdiagnostic samples compared to controls. Specifically, the SOM identified a distinct group of down-regulated miRNAs for lung cancer. The least pronounced group of down-regulated miRNAs was found for breast cancer. The most prominent group of higher expressed miRNAs was found in colon cancer and less prominent groups of down-regulated miRNAs for lung and breast cancer. In summary, the SOM analysis supports the presence of time- and disease specific miRNA patterns leading to the question which miRNAs contribute to these patterns.

### Diagnostic miRNAs

We computed for each miRNA the number of the above comparisons where it was higher- or lower expressed in any cancer or any time point compared to control samples (Fig. 5a). We found 59 miRNAs, which were not attributed to any time-point or disease. The majority of the miRNAs (222) was deregulated for only a single time point or a single cancer type. Of these, 90 (41.4%) were higher expressed and 130 were lower expressed (58.6%). We also observed miRNAs that were deregulated at most of the time points and for most cancer cases (top left and top right in Fig. 5a). The most prominent factor with the strongest impact on miRNA deregulation was the time point after cancer diagnosis as exemplified for miR-575 in Fig. 5b. However, we also found miRNAs that showed an increase of expression over time prior to diagnosis. An example of such a time course is miR-5006-5p in lung cancer as shown in Fig. 3c. Other miRNAs were lower expressed in pre-diagnostic cancer samples as compared to controls. An example is miR-6873-3p that showed an increase with age for the controls, but no comparable increase for the matched cancer samples (Fig. 5d/e). Since several time-points for the same individuals were measured we could generally ask whether the miRNA expression levels at consecutive time points were significantly altered in cancer cases but not in controls. A paired hypothesis test identified 14 miRNAs with significantly lower p-values for cases than for controls including miR-5196-5p and miR-320a as shown in Fig. 5f/g.

**Figure 5.**
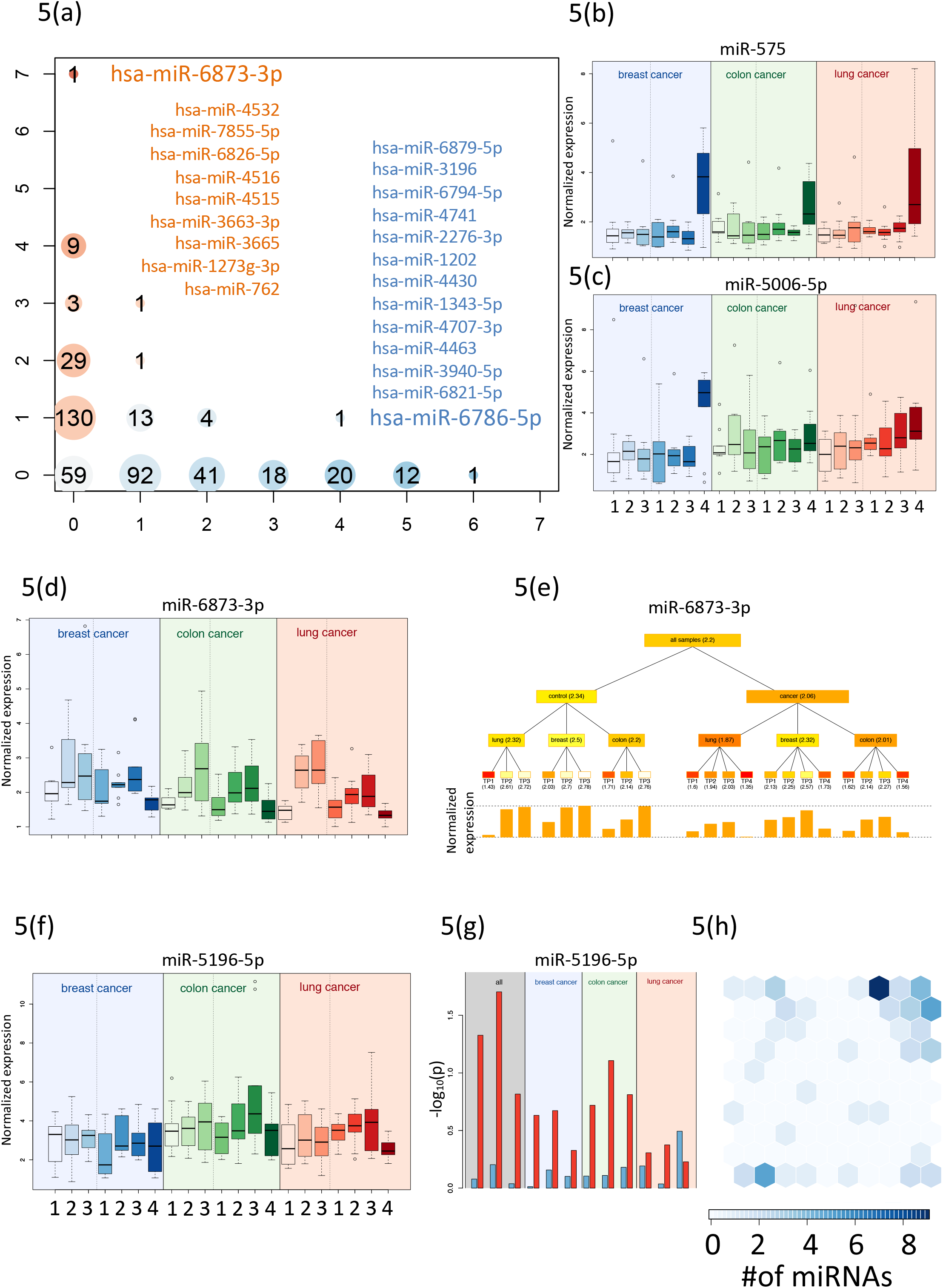
Specific miRNA patterns. (a) Computation of the number of significantly higher- or lower expressed miRNA for each time point and each cancer. Higher expressed miRNAs are indicated in red and lower expressed miRNAs in blue. The bubble size corresponds to the number of miRNAs found for a given time point and a specific cancer. Specifically indicated in blue is miR-6786 that were lower expressed in 6 analyses and higher expressed in none of the analyses. Also indicated in blue are the 12 miRNAs with lower expression in 6 analyses and higher expression in none of the analyses. Specifically indicated in orange is miR-6873-3p that were higher expressed in 7 analyses and lower expressed in none of the analyses. Also indicated in orange are the 9 miRNAs with higher expression in 4 analyses and lower expression in none of the analyses. (b) Box-plot of the normalized expression values of miR-575 for three time points prior to cancer diagnosis and one time point after diagnosis shown in the panels on the right side for each tumor type and the controls matched to the pre-diagnostic samples shown in the panel of the left side. The most prominent changes are found for the samples after diagnosis for all three cancer types. (c) Box-plot of the normalized expression values of miR-5006-5p presented as in Fig. 5b. MiR-5006-5p shows steadily increasing expression over time for lung cancer with the lowest expression value at the first time point and the highest expression at the latest time point. (d) Box-plot of the normalized expression values of miR-6873-3p presented as in Figure 5b. MiR-6873-3p shows a steadily increasing expression over time both for the controls and for the cancer samples. As also shown in Fig. 5e this increase is more prominent in the controls than in the cancer samples. The numbers represent the average normalized expression intensity. (e) Expression ontology. The expression for all groups is shown as tree structure. The leaves are the 21 groups of three cancers time seven total time points. Internal nodes contain the average of all nodes in the hierarchy below this node. The bar graph at the bottom represents the expression intensity of the leaves. (f) Box-plot of the normalized expression values of miR-5196-5p presented as in Fig. 5b. MiR-5196-5p shows a higher variability in the cancer samples than in the control samples. (g) Negative decade logarithm of paired t-test p-values for miR-5196-5p between cancer samples with all time-points combined and controls also with all time-points combined. In all cases the values for cancer indicated by red bars significantly exceed the values for the controls. (h) Allocation of 67 miRNAs, which were identified with a significantly altered expression by the different comparisons to the SOMs shown in Fig. 3 and 4. All miRNAs were identified by the SOM analysis that shown a strong enrichment for these miRNAs (upper right corner of the SOM map).

In total, we identified 93 miRNAs by the above 4 analyses including i) ANOVA of control samples, pre-diagnostic samples, and post-diagnostic cancer samples, ii) comparison of prediagnostic and controls samples, iii) identification of miRNAs that were deregulated in different cancer types, and iv) paired hypothesis test for miRNAs that are significantly altered in tumor samples but not in matched controls. Some miRNAs were identified by several analyses, reducing the number to 67 relevant miRNAs (Supplemental Table 3). The majority of these miRNAs have also been identified by the artificial intelligence-based analyses using SOMs. As shown in Fig. 5h SOM grouped 36 of these miRNAs in one cluster and 12 miRNAs in a second cluster. Notably, the miR-4687-3p and miR-6087 have been identified by three of analyses underlying their specific potential as pre-diagnostic markers.

## Discussion

Markers that facilitate detection of tumors in early stages, at best pre-diagnostic markers, are one of the most promising tools to improve cancer outcome. One question is how long prior to diagnosis molecular changes can be measured. To address such issues, very large sample collections are mandatory. Janus Serum Bank in Oslo (Norway) is an comprehensive resource for serum samples that have been stored and followed up over decades. A major strength of the Janus Serum Bank is the large number of collected samples and the long follow-up time, allowing to identify for each cancer case a matching control even by applying stringent criteria. The possibility for closely matched pairs was essential for the design of our study (nested case-control design), which analyzed 30 pairs of samples, each including an individual who developed cancer and a matched individual who was not diagnosed with cancer at any time point. A further strength of the Janus Serum Bank is the multiple sampling over time allowing to follow up pre-diagnostic changes of a biomarker. This characteristic was also central to the design of our study, which includes three pre-diagnostic samples of each individual selected. One limitation in the study set up was a later time point for controls. Since the collection to the Janus Serum Bank was ended in 2004 we were not able to acquire matched controls for the diagnostic samples, which were obtained at later time points.

Already in previous research studies we obtained valuable results from samples of the Janus Serum Bank. We determined the influence of confounding factors including storage time, age, sex, smoking, and body mass index among others on the patterns ^18^ of blood-borne miRNAs. Further, we identified pre-diagnostic miRNA patterns in sera from lung cancer patients ^19^. In the present study we extend our previous results with respect to many aspects. We now include more samples per cancer type, we include time points that are much further away from the diagnosis and we measure and compared different cancer types. Lastly, we matched on the time difference between the samples allowing only minimal variation (a maximal difference between the blood collection time point of cases and controls of only up to two months). The extended scope of the study also called for different bioinformatic and biostatistical approaches, such as self-organizing maps as an artificial neural network approach.

The different design of our present and our previous analysis makes it difficult to compare the two analyses both in terms of sample collections and in terms of the applied measurement technology. Despite the substantial differences between the studies, we reidentified several pre-diagnostic miRNAs from the previous study in our present results, including miR-762, miR-1202, miR-1207-5p, and miR-575. To further gauge the biological meaning and especially the evidence for causative role of the pre-diagnostic miRNAs identified in the present study, we evaluated these miRNAs with regards to their previously reported involvement in cancer. A systematic PubMed search for the 67 miRNAs identified in our study yielded 324 manuscripts that report a cancer connection for these miRNAs (Supplemental Table 4). Most frequently we found a cancer association for miR-320a, which was described in 93 studies, for miR-630 in 59 studies, for miR-1207-5p in 25 studies, and for miR-149-3p in 21 studies. Specifically, miR-320a has been associated with lung carcinoma in 11 studies, with breast carcinoma in 17 studies and with colon carcinoma in 8 studies. Likewise, miR-630 and miR-1207 have previously been associated with the cancer types analyzed in the present study. In detail, miR-630 was associated in 10 studies with lung cancer, in 6 studies with breast cancer, in 2 studies with colon cancer, and miR-1207-5p in 4 studies with lung cancer, in 4 studies with breast cancer and in 2 studies with colon cancer (Supplemental Table 5). Our results emphasized miR-4687-3p to have specific potential as pre-diagnostic marker. This is also suggested by Nagy et al who identified altered levels of miR-4687 in plasma of colorectal cancer and adenomas cases compared to individuals with normal colon ^30^. Also miR-6087, the other miRNA showing specific potential, has previously been suggested as a circulating early detection biomarker of bladder cancer in a large serum study ^31^, with a potential role in regulation of p53 ^32^. Circulating cell-free RNA are considered promising as liquid biopsy cancer markers, although the biological and clinical interpretations are challenging ^33^. Our study is a indication of RNA profiles specific for cancer decades before diagnosis, however, validation and biological and clinical interpretations are needed in future studies.

Since most of these studies were on tissue or cell cultures, care must be taken to prematurely hypothesize a causal role for the pre-diagnostic sera miRNAs in the development of these cancers.. Although we analyzed a total of 210 samples, it has also to be acknowledged that we could only identify 10 patients for each cancer type. This was due to the criteria of sample selection requiring i) three pre-diagnostic samples for each patient with ii) comparable collection time points, and with iii) matched controls both in terms of the patients’ age and the collection time points. The predictive value of the identified miRNAs awaits confirmation by prospective studies with an extended number of samples ideally recruiting from additional populations beside the Norwegian population that was analyzed in the present study. Such prospective cohort-based studies, however, require long follow up times as part of a longitudinal study design, which are essential to identify new biomarkers ^34^.

In summary, our results suggest that circulating miRNA signatures can be found decades prior to the clinical manifestation of a tumor. The most prominent miRNA changes occur in pre-diagnostic samples for lung cancer, which could however be confounded by smoking behaviour of patients and controls. This is consistent with our previous study that showed dynamic pre-diagnostic changes of circulating RNAs related to the histology and the stage of lung cancer after its manifestation. As for colon and breast cancer, our results indicate less pronounced changes of blood-borne miRNAs prior to diagnosis. While the results of our study are generally promising it is evident that reproduction in other cohorts is required.

## Methods

### Study Set-up and RNA extraction

In the study we included lung cancer, colon cancer and breast cancer patients, three of the top most common cancers and where the identification of early detection biomarkers would have a large impact. We also wanted to compare miRNA signatures across different cancer types. The cancer cases were identified by linking the Janus Cohort to the Cancer Registry of Norway using the individual’s Norwegian national identity number.

For each cancer group, 10 patients were included who had three pre-diagnostic and one diagnostic sample available. 30 control individuals were selected. For the cancer patients three pre-diagnostic and one post diagnostic time point were measured (120 samples) for the controls three time points matching the pre-diagnostic time points were measured (90 samples) (figure 1 (b)). Between cancer and matched controls, a maximal time difference of two months was allowed. All samples in the JSB are stored at −25°C and collected in gel vials, or in 10-mL tubes containing either 5 mg sodium iodoacetate or no additives. Total RNA including miRNAs was isolated using the miRNeasy Serum/Plasma Kit (Qiagen, Hilden, Germany) as previously described.^17^ Of the 210 samples, 204 yielded high-quality RNA and microarray results, 6 samples were excluded for quality reasons.^35^ The study was approved by the Norwegian regional committee for medical and health research ethics (REC no: 2013/614). The donors have given broad consent for the use of the samples in cancer research.

### Microarray measurement

Genome wide miRNA expression profiles were created using the SurePrint G3 8 × 60k miRNA microarray (miRBase version 21, Cat. no. G4872A). Using this microarray, probes for 2,549 mature human miRNAs were measured. As input material for the microarray screening, 100 ng total RNA including miRNA was used for each sample. The hybridization process and read out of the microarrays has been performed according to manufacturer’s recommendations as previously described.^17^

### Data processing and bioinformatics

Features were extracted from the manufacturers GW Feature Extraction software (version 10.10.11, Agilent Technologies). Replicated measurements of miRNAs were summarized by the median expression and data were subjected to standard quantile normalization. Filtering of miRNAs close to the background excluded 2,114 miRNAs leaving an expressed set of 435 serum miRNAs. The filtering was done using the present call definition of the Manufacturers software that identifies a feature to be expressed if it is significantly above the microarray background. Since miRNA measurements were not always normally distributed (according to Shapiro Wilk Normality tests), non-parametric Wilcoxon Mann-Whitney (WMW) test have been performed in addition to the parametric t-test. If not mentioned explicitly, p-values in the manuscript rely on the WMW test. Because of the explorative nature of our study nominal p-values are reported. To assess differential expression of miRNAs, the area under the curve (AUC) has been computed in addition to p-values. Here, an AUC close to 0.5 means no dys-regulation, an AUC close to 0 means higher expression in controls and an AUC close to 1 means higher expression in cancer patients. To compute the density of AUC values, the em algorithm for mixtures of univariate normal the normalmixEM from the R mixtools package has been applied assuming two components. To assess sources of data variability, Principal Variance Component Analysis (PVCA) using the Bioconductor pvca package has been performed. To learn cancer and time patterns we applied one type of artificial neural networks (aNN), so called self-organized maps (SOMs). The computations have been performed using the kohonen and somgrid package from R. We used a hexagonal 10×10 grid to group the 435 serum miRNAs. As feature vector, the AUC values for the different comparisons were used. The data set was presented 10,000 times to the SOM with a learning rate linearly decreasing from 0.05 to 0.01. To cluster the SOM results, hierarchical clustering using the hclust function has been performed.

